# Sphingolipid Containing Outer Membrane Vesicles Serve as a Delivery Vehicle to Limit Host Immune Response to *Porphyromonas gingivalis*

**DOI:** 10.1101/2020.10.05.327544

**Authors:** Fernanda G. Rocha, Gregory Ottenberg, Zavier G. Eure, Mary E. Davey, Frank C. Gibson

## Abstract

Sphingolipids (SLs) are essential structural components of mammalian cell membranes. Our group recently determined that the oral anaerobe *Porphyromonas gingivalis* delivers its SLs to host cells, and that the ability of *P. gingivalis* to synthesize SLs limits the elicited host inflammatory response during cellular infection. As *P. gingivalis* robustly produces outer membrane vesicles (OMVs), we hypothesized that OMVs serve as a delivery vehicle for SLs, that the SL status of the OMVs may impact cargo loading to OMVs, and that SL-containing OMVs limit elicited host inflammation similar to that observed by direct bacterial challenge. Transwell cell culture experiments determined that in comparison to the parent strain W83, the SL-null mutant elicited a hyper-inflammatory immune response from THP-1 macrophage-like cells with elevated TNF-α, IL-1β, and IL-6. Targeted assessment of Toll-like receptors (TLRs) identified elevated expression of TLR2, unchanged TLR4, and elevated expression of the adaptor molecules MyD88 and TRIF by SL-null *P. gingivalis.* No significant differences in gingipain activity were observed in our infection models and both strains produced OMVs of similar size. Using comparative 2-dimensional gel electrophoresis we identified differences in the protein cargo of the OMVs between parent and SL-null strain. Importantly, use of purified OMVs recapitulated the cellular inflammatory response observed in the transwell system with whole bacteria. These findings provide new insights into the role of SLs in *P. gingivalis* OMV cargo assembly and expand our understanding of SL-OMVs as bacterial structures that modulate the host inflammatory response.

## INTRODUCTION

Microbial outer membrane vesicles (OMVs), also referred to extracellular vesicles are spherical nanosized proteoliposomes that are generated by vesiculation of the bacterial outer membrane. OMVs have been shown to represent a key mode of interkingdom communication between bacteria and host tissues (1). The size of OMVs range from 20 to 400 nm, and they are comprised of a lipid bilayer containing lipopolysaccharide, outer membrane proteins; and other bioactive effector molecules such as periplasmic and cytosolic proteins, nucleic acids, and immunomodulatory factors (2, 3). The mechanism of OMV production is partially understood (1). It is also known that the biogenesis of OMVs is beneficial for bacterial pathogens, as these structures are known to deliver multiple virulence factors to the host. In addition, OMVs concentrate and protect virulence determinants from host degradative process such as proteases, confer antibiotic resistance, and facilitate biofilm formation (2, 4). Several studies have demonstrated that purified OMVs can enhance microbial pathogenicity by triggering the release of proinflammatory cytokines, as well as promoting neutrophil migration and disruption of epithelial cell junctions (5, 6). Not surprisingly, some beneficial effects have been linked to OMV production by bacteria from the host gut microbiota (4).

Periodontitis is among the most common bacteria-elicited chronic inflammatory diseases of humans (7). In susceptible individuals, bacteria in the subgingival biofilm promote dysregulated inflammation and microbial dysbiosis that results in the progressive destruction of soft and hard tissues that support the teeth, which is irreversible (8). Locally, one of the characteristics of sites affected by periodontitis is the increase of Gram-negative anaerobes, such as *Porphyromonas gingivalis, Tannerella forsythia,* and *Treponema denticola,* that are abundant in specific virulence factors, including gingipains, lipopolysaccharide, capsular polysaccharide, lipoproteins, and other molecules (9, 10). Interestingly, several bacteria associated with periodontal disease including *P. gingivalis,* are known OMV producers (6, 11, 12).

There is significant interest as to how *P. gingivalis* evolves and contributes to the changing inflammatory milieu from periodontal health to disease. Indeed, although this organism is closely associated with periodontal disease, this bacterium is found in subgingival plaque during periodontal health (13), suggesting that the host does not always responds to *P. gingivalis* as a pathogen. An area of intense focus has been the ability of *P. gingivalis* to perform targeted immunosuppression and immune evasion by deploying molecules including its gingipains, atypical LPS, and other factors to disrupt the host immune response (14–16). Recently, our group reported that *P. gingivalis* possess an additional layer of immunosuppressive ability that is mediated by its ability to synthesize sphingolipids (SLs). Furthermore, and intriguingly, the SLs are delivered to host cells in a contact-independent manner (17). Considering that *P. gingivalis* is known to be a robust OMV producer, we here hypothesized that SL-containing OMVs (SL-OMVs) may serve as a unique delivery platform, and that the SL-OMVs can limit the host immune response.

In this study, we purified OMVs from SL containing or SL-null *P. gingivalis* and assessed the protein content of these vesicles. In addition, using cell culture, we characterized the immunomodulatory effect of whether *P. gingivalis* synthesizes SLs on the host immune response. Moreover, we demonstrate that in pure form SL-OMVs and OMVs lacking SLs recapitulate the host immune response observed in response to parent and mutant *P. gingivalis* infection.

## RESULTS

### SL synthesis limits the inflammatory response of immune cells to *P. gingivalis* in a contact-independent manner

To investigate the immunoregulatory effect of *P. gingivalis* SL synthesis in cell culture without direct contact with host cells, we utilized a 0.4 μm-pore transwell system, hence the OMVs (<400 nm) can cross the transwell membrane, while preventing direct bacterial-host cell contact. We observed that stimulation with SPT-mutant elicited a more robust immune response from THP-1 cells compared to the response elicited by the WT (Fig. 1A). As early as 2 hrs after initiation of transwell co-culture, significantly higher levels of TNF-α were measured in supernatant fluids from THP-1 cells cultured with SL-null *P. gingivalis* compared with parent (p < 0.05; Fig. 1A), a trend that was maintained through the 24 hrs infection period. There was also a significant increase in the levels of IL-1β and IL-8 by 6 hrs elicited by SL-null mutant (p < 0.05 for all by t test). The trend of hyperinflammatory response when SLs were not synthesized could also be observed in the measurements of IL-6, IL-10 and RANTES (Fig. 1A). These data support the concept that SL production limits the inflammatory response of immune cells to *P. gingivalis,* and that this immunoregulatory capacity does not require bacteria-to-host cell contact.

**Figure 1.**
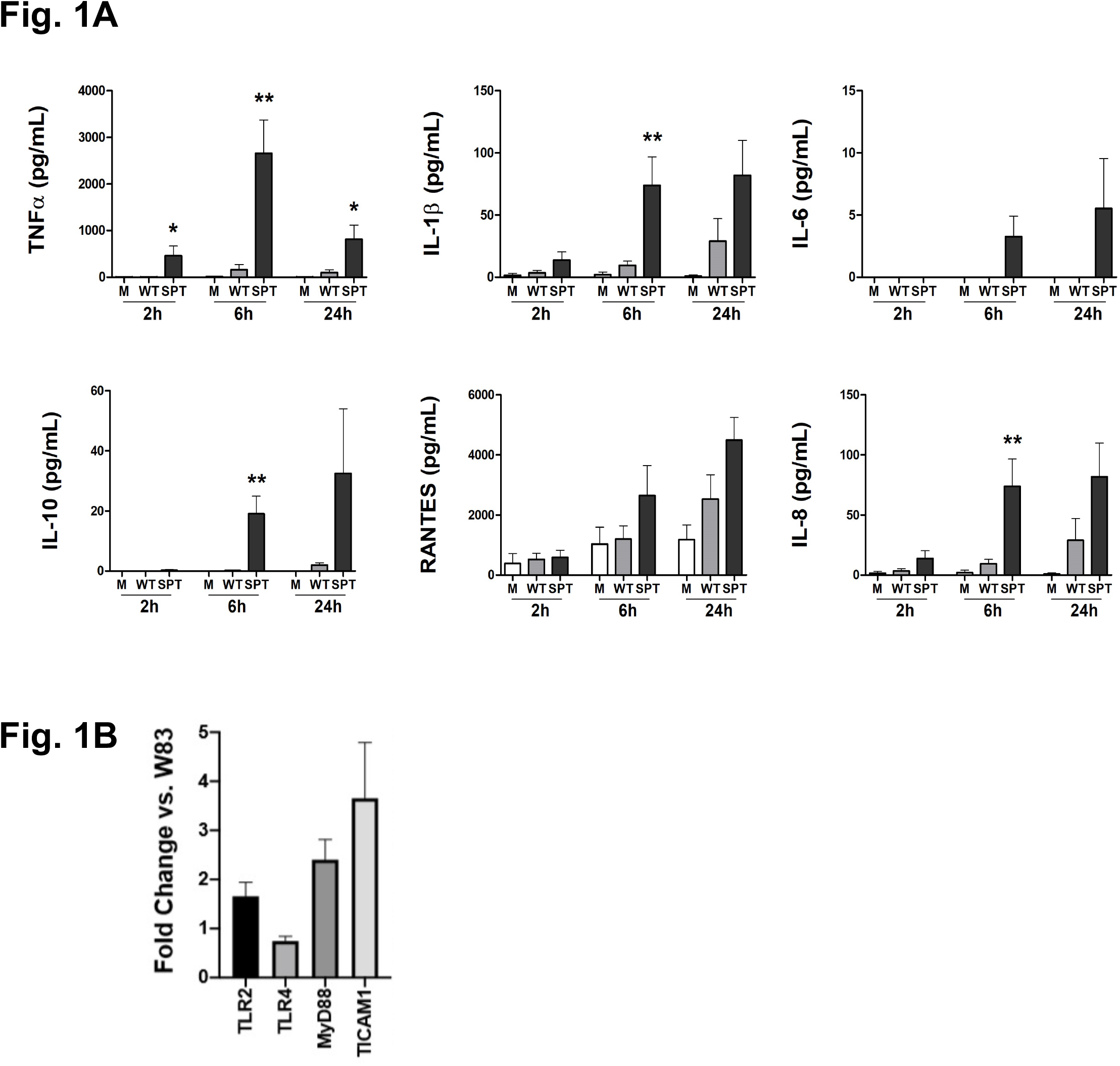
*P. gingivalis* sphingolipids suppress host immune response independently of direct contact. Supernatant fluids were collected from PMA-activated human THP-1 cells placed in the lower well of a transwell system and to the upper well was added either *P. gingivalis* W83 (WT; gray bars) or the *P. gingivalis* W83 SL-null mutant (SPT-; black bars) at 2, 6 and 24 hrs of culture, and the levels of TNF-α, IL-1β, IL-6, IL-10, RANTES, and IL-8 were measured by multiplex immunoassay (**A.**). Medium alone (M; white bars) served as unchallenged control. (**B.**) Transcriptional levels of TLR pathway genes from THP-1 cells cultured in transwells, with medium, *P. gingivalis* W83 WT, or *P. gingivalis* SPT-for 2h. Expression data were normalized to β-actin, and fold-change in TRL pathway genes are presented for SPT relative to W83. Data are presented as mean +/-SEM (n = 3 independent experiments); * = P <0.05 and ** = P <0.01 vs. WT *P. gingivalis* using unpaired t-tests.

Considering the differences in the cytokines and chemokine levels in the THP-1 cells supernatants, understanding that some *P. gingivalis* SLs are sensed by TLRs (18), and the knowledge that OMV activation of host cells is mediated in part by TLRs (19), we subsequently evaluated gene expression of selected TLR pathway members: TLR2, TLR4, MyD88, and TICAM/TRIF. In comparison to WT *P. gingivalis,* stimulation with the SPT-mutant resulted in increase of TLR2 gene expression (p < 0.05) without notable change in the expression of TLR4 (Fig. 1B). Investigating the expression of the key adaptor molecules responsible for TRL signaling, we observed increased expression of MyD88 and TICAM1/TRIF when THP-1 cells were stimulated with SPT mutant, in comparison to WT (p <0.05 and p <0.05, respectively; Fig. 2B). These data support that when *P. gingivalis* can synthesize SLs, this limits the ability of the host cell to mount a robust immune response to this organism. Moreover, since the regulation occurred in a transwell system, the most likely delivery mechanism is via SL-containing OMVs.

**Figure 2.**
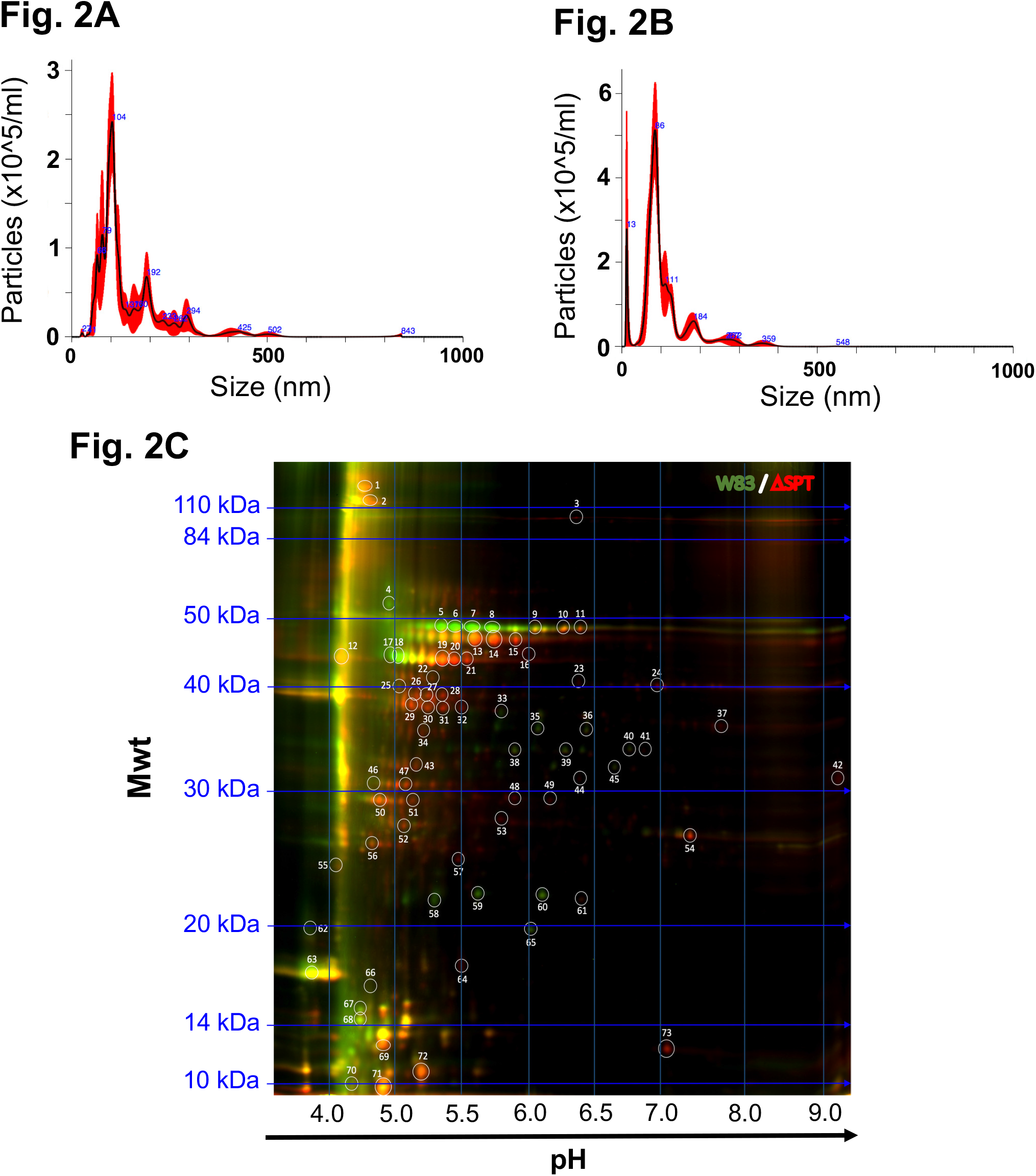
Characterization of *P. gingivalis* OMVs. NanoSight analysis of OMV from *P. gingivalis* W83 WT (**A.**) and *P. gingivalis* ΔSPT mutant (**B.**). (**C.**) Comparative fluorescent two-dimensional gel of OMV protein cargo from *P. gingivalis* W83 WT (green spots) and the ΔSPT mutant (red spots). Proteins common in OMVs of the WT and ΔSPT strains are yellow spots.

### Nanosight characterization of *P. gingivalis* OMVs from SL-producing (WT) and SL-null mutant (SPT) and protein profiles

OMVs were isolated from the culture medium of the WT and SPT-mutant and analyzed using NanoSight, and the size distribution was comparable between WT and the SPT-mutant (Fig. 2A). Further, the SPT-mutant was found to be more proficient than WT in OMV production (Fig. 2B), suggesting that the inability to synthesize SLs may lead *P. gingivalis* to increase the production of other types of OMVs.

To begin to understand how SL synthesis may influence *P. gingivalis* OMV cargo, comparative 2-dimentional gel electrophoresis was used to determine the proteins present in purified OMVs isolated from the WT and the SPT-mutant. Remarkably, several proteins detected in the SL-containing OMVs were absent in the SL-null OMVs (Fig. 2C). One dominant protein was identified as PG1881, a predicted pilin lipoprotein recently reported to be central to OMV production, and potentially linked to vesiculation mechanisms (20). Such differences in the protein content support our hypothesis that like lipid rafts in eukaryotes (21), SL-containing microdomains in the outer membrane of *P. gingivalis* contain select proteins, which are ultimately packaged in SL-OMVs, thereby contributing to the ability of SL-OMVs to be immunosuppressive.

### Immunomodulatory effect of *P. gingivalis* SL synthesis occurs without interference of active gingipains

Gingipain activity is known to modulate the ability of the host to mount a proper immune response by degrading immunologically important molecules (22), therefore we assessed the possibility of interference of active gingipains in our assay. Gingipain assays (both arginine- and lysine-specific) performed on THP-1 cell culture supernatant fluids showed only nominal gingipain activity without addition of a reductant (normoxic conditions), and similar levels of activity were detected in the cell culture supernatant fluids of WT and SPT-mutant when tested under typical reducing conditions (10 mM cysteine) as a control (Supplemental Data Fig. 1). Inspection of gingipain activity from bacterial culture supernatant fluids and cell pellets from WT and SPT-*P. gingivalis* revealed that the culture supernates of *P. gingivalis* SPT-mutant possessed slightly higher, not lower levels of gingipain activity as compared with WT (Supplemental Data Fig. 1), while similar levels of gingipain activity were observed between WT and SPT-mutant cells pellets (Supplemental Data Fig. 1). Our findings show that the observed low-level inflammation observed in response to WT *P. gingivalis* in the transwell experiments does not align with elevated gingipain activity.

### Purified OMVs suppress the host cell inflammatory response

To evaluate the possibility that purified *P. gingivalis* OMVs are able to recapitulate the immune stimulating activity previously observed by direct bacterial challenge (17), and no-contact / transwell challenge, purified OMVs from WT and SPT-mutant were added directly to THP-1 cells. We detected a hyperinflammatory response from THP-1 cells cultured directly with OMVs from the SL-null mutant in comparison to the elicited response by the OMVs from WT (Fig. 3). The levels of TNF-α and IL-1β were significantly higher at 2 hrs and 6 hrs after stimuli with SLs-null mutant than WT (p < 0.05 for all by t test), and that same trend was evident at 24 hrs. The OMVs produced by the SL-null strain also elicited higher levels of IL-6, IL-8, RANTES and IL-10, with a significant increase observed at 24 hrs (Fig. 3). These findings confirm that purified *P. gingivalis* OMVs recapitulate the findings for suppression of the immune response to *P. gingivalis* observed when we evaluated systems using whole live bacteria.

**Figure 3.**
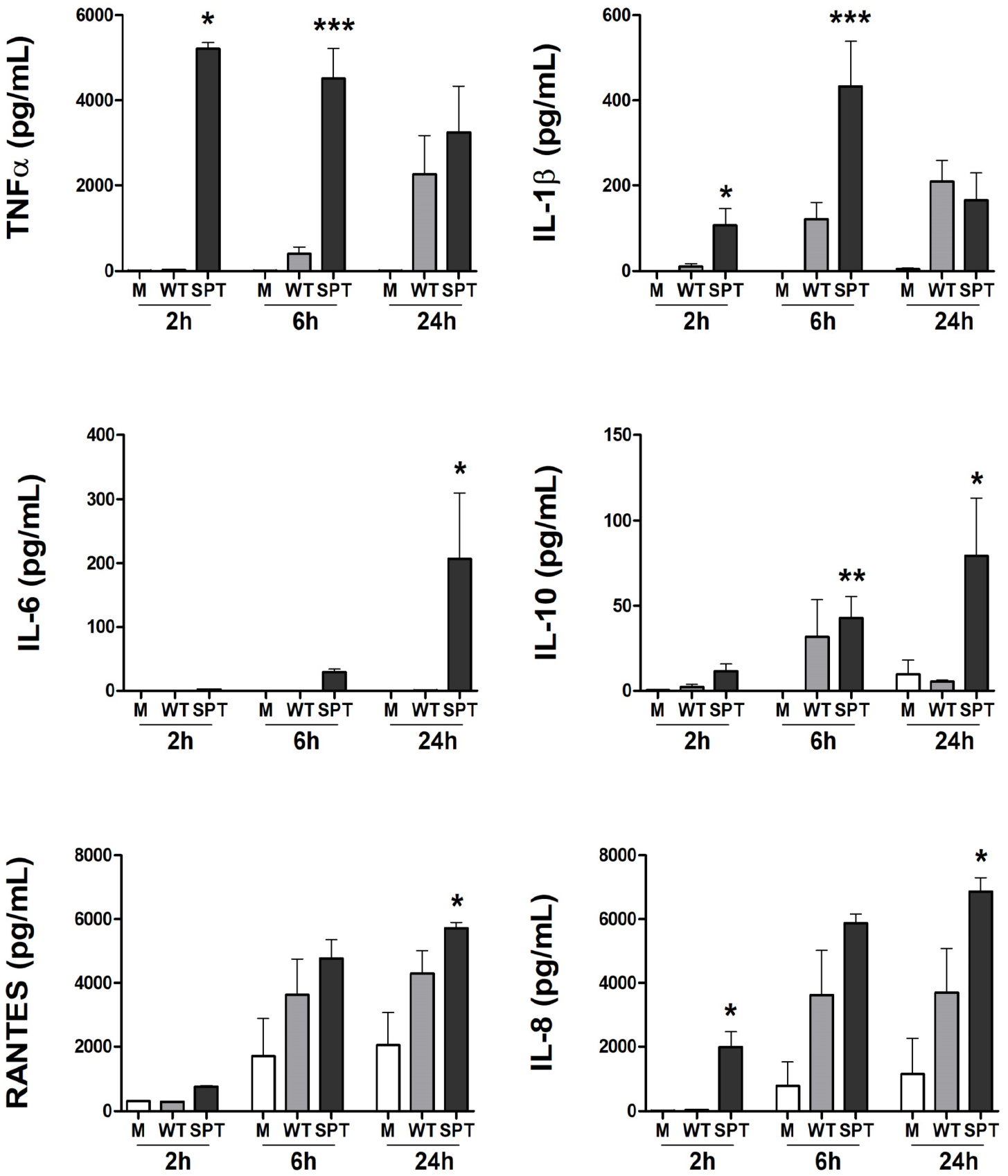
Purified OMVs from *P. gingivalis* from SLs null mutant elicited higher cytokine and chemokine response. PMA-treated human macrophage-like THP-1 cells were directly cultured with purified OMVs (1000 particles / cell) from *P. gingivalis* W83 (WT; gray bars) or the *P. gingivalis* W83 SL-null mutant (SPT-; black bars). Supernatant fluids were collected at 2, 6 and 24h, and the levels of TNF-α, IL-1β, IL-6, IL-10, RANTES, and IL-8 were measured by multiplex immunoassay. Medium alone (M; white bars) served as unchallenged control. Data are presented as mean +/− SEM (n = 4 independent experiments); * = P <0.05, ** = P <0.01, and *** = P <0.005 vs. WT *P. gingivalis* using unpaired t-tests.

## DISCUSSION

The concept that *P. gingivalis* SLs participate in modulating host inflammation during infection is a critical point of novel understanding that is further supported and extended in the present study. As *P. gingivalis* SLs have been recently shown to be transferred to host cells, and this transfer correlates with suppression of the inflammatory response from those cells (17), we speculated that SL-OMVs act as a delivery platform for transfer. It is understood that *P. gingivalis* is a robust producer of OMVs, that these nanostructures can elicit inflammatory responses (3), and the mechanism of elicited inflammation is in part dependent on the innate immune sensing of these particles by host cells (6, 19). However, there is still a gap in the knowledge of the role of OMVs from the different type strains of *P. gingivalis* and in inflammatory immune cells, such as macrophages. Cecil *et al.* (6) found that OMVs from *P. gingivalis* strain W50 strongly interacted with monocytes and macrophages, increasing their pro-inflammatory responses, in a dose-dependent manner. Lower concentrations of OMVs induced stronger immune response, while higher concentrations were less inflammatory in unprimed cells, but stronger upon second exposure. OMVs from *P. gingivalis* strain W50 have also shown to induce a metabolic shift in macrophages, with strong activation of proinflammatory cytokine production, inflammasome signaling, and pyroptotic cell death, in contrast to a weak cytokine production, with no inflammasome activation or pyroptosis induction by macrophages that were stimulated with the whole bacteria (23). In this study, we demonstrated that macrophages cultured with purified OMVs from *P. gingivalis* strain W83 produced a mild inflammatory phenotype consistent with live bacterial challenge, while OMVs from SPT-mutant were found consistent with a hyper-inflammatory phenotype when the SLs were absent. More studies are required to obtain a better understanding of how different strains of *P. gingivalis* impact the interaction between OMV and host cells.

The fact that OMVs deliver *P. gingivalis* SLs to the host cell is particularly fascinating when we consider the periodontal environment, since previous studies uncovered that microbial SLs are present at differing levels in gingival tissues of healthy individuals and patients with periodontal diseases (24). Intriguingly, bacterial SLs seem to go through a shift in their dihydroceramide (DHC) pools that matches the transition from health to disease. In the diseased gingival tissues, the lipid pools present low phosphoglycerol-DHC (PG-DHC) lipid levels and an increase in phosphoethanolamine-DHC (PE-DHC) lipids (24). These findings suggest that synthesis of different SLs by the oral Bacteroidetes may influence periodontal homeostasis as well as disease progression, thus, the different SL pools may be related to both conditions of the gingival tissues. *In vitro* studies using isolated *P. gingivalis* SLs have shown effects of purified individual SLs in bone remodeling and inflammatory expression in some cells (18, 25–30), and our recently published data, utilizing the whole bacteria. These findings support the fundamental role of *P. gingivalis* SLs in immune-regulation (17); suggesting that SLs have multiple functions. Future research, including studies aimed at defining SL impacts on other immunologically important cell types including neutrophils, and others, as well as animal modeling is needed to decipher the importance of specific *P. gingivalis* SLs in the context of infection-elicited host response and linkage to periodontal disease.

In an earlier study, we determined that WT *P. gingivalis* OMVs contain SLs and the STP-mutant was devoid of SLs (31). Here, we explored the impact of SL synthesis on the characteristics of OMVs produced by *P. gingivalis* by comparing OMVs from the SL producing WT and SL-null mutant. The OMV size distribution that we observed is comparable to that detected by others, which was reportedly to range from 50 to 300 nm in diameter (19, 32). We observed that *P. gingivalis* produced OMVs even when it cannot synthesize SLs, and that OMV production was greater in the SL mutant than WT, suggesting that SL synthesis influences biogenesis of OMVs that do not contain SLs, or that SL synthesis negatively regulates production of other OMVs that do not possess SLs. Like other bacterial OMVs, the OMVs of *P. gingivalis* are known to possess an array of molecules including proteins and LPS, as well as its peptidoglycan (33). To begin to understand the contribution of SLs to the loading of cargo to purified OMVs, we compared the protein profiles between wild type and SL-null OMVs and noted some dramatic differences, indicating that SL-synthesis impacts OMV cargo. Recent publications have shown that proteins carried on OMVs produced by *P. gingivalis* strain W83 are citrullinated by PPAD and this included protein PG1881 (20). The data presented in this study demonstrate that when *P. gingivalis* cannot synthesize SLs, it was proficient in OMV biogenesis, yet the PG1881 protein is absent in the OMVs. Since, as noted, we previously determined that *P. gingivalis* OMVs can contain SLs (31), this finding suggests that SL-containing OMVs are enriched in PG1881.

Importantly, a recent report by (34) showed that PG1881 is a predicted pili forming lipoprotein that plays a role in vesiculation. Thus, our working model is that certain subtypes of SLs form membrane microdomains which directs localization of PG1881 to these SL-rich areas. Studies are on-going to elucidate the mechanisms involved in SL-OMV biogenesis.

As previously demonstrated, the direct challenge with the SLs null mutant elicited a hyperinflammatory profile from the host cells (17). Here, a highly similar cytokine profile was detected from macrophages cultured with wild type and SL-deficient *P. gingivalis* when bacteria to the host cell contact was prevented, by using a transwell system (Fig. 4). The inflammatory profile detected in this study using wild type *P. gingivalis* has a close similarity to that found in periodontal disease, with elevated levels of pro-inflammatory cytokines TNF-α, IL-1β and IL-6 and chemokines such IL-8, which are characteristically found in gingival tissues of periodontal disease patients (35), and induce higher levels of IL-10, an anti-inflammatory cytokine that has been showed to be upregulated in gingival crevicular fluid of periodontal patients and known to suppress the production of pro-inflammatory cytokines from several cell types (36, 37). The inflammatory response was exacerbated by absence of SLs in a contact-independent manner, suggesting that SLs containing OMVs are capable of regulating inflammation, and this ability was impacted by the different cargo loading of the SL-null OMVs, consequently, eliciting a hyper inflammatory profile. The mechanisms by which this occurs are beginning to be explored. Interestingly, our findings are consistent with other closely related bacteria such as *Bacteroides thetaiotaomicron* in the intestinal tract, where *B. thetaiotaomicron* SL production and uptake by host tissues is important to maintain gut homeostasis as a SL-deficient mutant elicited significantly higher amounts of inflammation in an inflammatory bowel diseases model (30, 38). Although it is not clear how *P. gingivalis* OMVs deliver their cargo to the host, we demonstrate here that purified OMVs from SL-null mutant elicit a stronger inflammatory response compared to OMVs from wild type *P. gingivalis.* These findings produce the same inflammatory profile we detected when using the whole bacteria, by both direct challenge and in the contact prevented transwell system. Noteworthy, SL containing OMVs can limit the inflammatory response of the host like the whole bacteria, indicating the fundamental role of *P. gingivalis* OMVs cargo in its interaction with the host cells. Strong interaction of OMVs from *P. gingivalis* with THP-1 macrophages has been reported, including induction of phagocytosis, NF-κB activation, cellular priming, and accentuated pro-inflammatory response (6). It is also well understood that OMVs can penetrate and disseminate through host tissues more easily than their larger parent bacterial cells, due to their nanoparticle size, proteolytic and adhesive properties (39).

**Figure 4.**
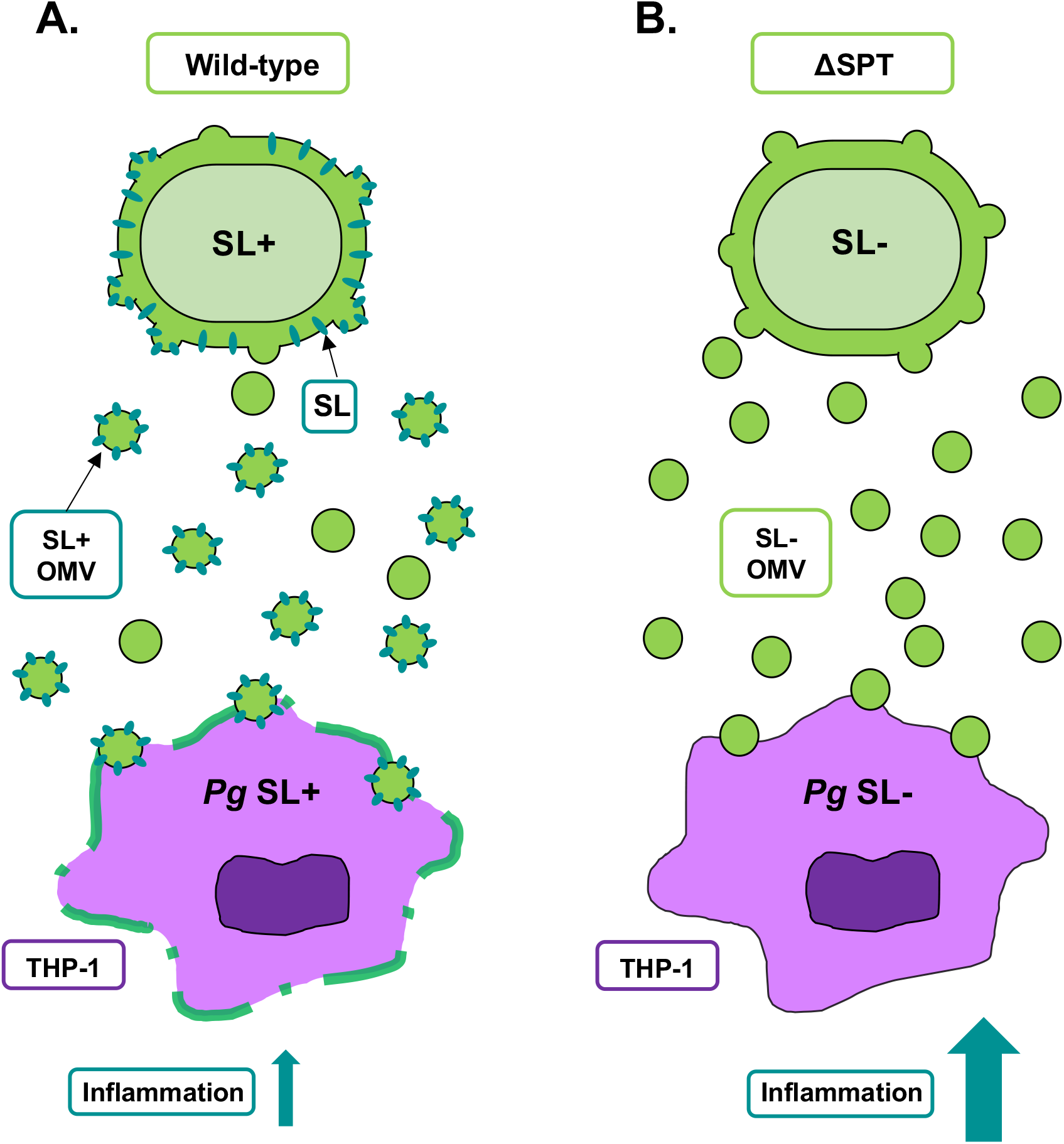
Proposed working model elucidating the role of *P. gingivalis* OMVs as a vehicle to deliver bacterial sphingolipids (SL) to host cells. In a transwell system, wild-type *P. gingivalis* are able to deliver its SLs to THP-1 cells in a contact independent manner (A). The SL-null mutant (ΔSPT) still produces OMVs (B), but as SLs are not synthesized, it elicits a hyper inflammatory response by the host cells in comparison to the wild-type, demonstrating the participation of *P. gingivalis* SLs in the regulation of inflammatory response by the host cells.

Our past work, and current studies have shown that TLRs (TLR2, TLR4, and others) and their signaling pathways are important in host innate immune sensing of *P. gingivalis* (40–43). Comparing the THP-1 cell response to SL-containing and SL-deficient *P. gingivalis* in the transwell system, we found that THP-1 cells cultured with SL-containing *P. gingivalis* presented reduced expression of TLR2 gene and TLR adaptor molecules MyD88 and TICAM1/TRIF as compared with the hyper-inflammatory SL-deficient mutant; however no differences in TLR4 gene expression were observed. Both TLR2 and TLR4 are critical receptors to control the induction of TNF-α, IL-1β, IL-6 and IL-8 by pathogens in the periodontal tissues (44), to *P. gingivalis* by cells exposed to the organism *in vitro* (41, 42, 45, 46), and it is also known that *P. gingivalis* OMVs utilizes TLRs *in vitro* (19). It is well established that TLR2 and TLR4 and their signaling pathways (MyD88 and TRIF) play an important role in how *P. gingivalis* is immunologically sensed by the host, and purified SLs have also been stated to signal through TLR2 (47, 48). In addition, it has been suggested in a study using purified *P. gingivalis* OMVs that these particles may be critical to inflammasome activation and stimulation of multiple PPRs, which may have a synergistic effect, perhaps even greater than the stimulation of TLR4 alone (19). Thus, further studies are necessary to obtain a better understanding of the mechanisms that control the immunosuppressive effect we observe in the presence of *P. gingivalis* SLs.

Although several pathways exist by which *P. gingivalis* can promote immune dysregulation (49), a well-established mechanism utilized by *P. gingivalis* is the degradation of many host proteins by gingipains, which are cysteine proteinases (50). Considered major virulence determinants of *P. gingivalis,* the arginine and lysine gingipains can be released from the organism via OMVs or directly secreted. Previous work from our group identified that the SL-null mutant produces lower cell-associated arginine and lysine gingipain activity when compared to the WT, with an increase in the release of gingipains into the supernatant (31). As we noted that there are protein cargo differences of OMVs from WT and SL-*P. gingivalis,* we also examined gingipain activity, both in conditions resembling the cell culture environment (normoxia) and using classical highly reducing conditions. Interestingly, almost no gingipain activity was detected in the cell culture supernates when THP-1 was cultured with wild type or with SL-null mutant in a transwell system. These findings showed that under cell culture conditions gingipain activity was essentially non-existent; however, when these same samples were examined under a reducing environment of traditional assay conditions, low levels of gingipain activity could be detected. Importantly, there was no difference in activity between WT and SL-mutant transwell supernatant fluids. In parallel, bacterial cell culture supernatant fluids and cell pellets grown in the same medium (RPMI) were used to validate the assay. The SPT mutant supernate possessed higher levels of gingipain activity when compared with WT *P. gingivalis,* as previous shown when the strains were grown in TSBHK (31). The importance of these data are that the supernatant fluids, where OMVs would be enriched, do not have more activity in the SPT mutant than the WT. Although this does not eliminate a role for gingipains, the findings indicate that a reduction in gingipain activity is not a likely explanation for the observed elevated levels of inflammatory molecules in response to *P. gingivalis* when the organism is unable to synthesize SLs. Our data, therefore, suggest that the mechanism of immunomodulation is mediated directly by the SLs on the OMVs or the cargo of the OMVs, or both; however the underlying mechanism(s) remain to be determined.

In the context of periodontal infection, OMVs can disrupt epithelial cells junctions and deliver virulence factors from bacteria to immune cells that are in the underlying tissues. Therefore, the concentration gradient of OMVs is likely to occur from the site of bacterial infection in the polymicrobial biofilm adhered to the tooth root. OMV concentrations then will tend to be lower at distal areas from the initial site of infection, priming the host cells, but not necessarily yet promoting cell activation. As the disease progresses, the tissues previously primed by exposure to OMVs will consequently have a more robust inflammatory response (6). Considering that these pro-inflammatory interactions depend upon early OMV interactions with macrophages and other immune cells, our data reveals that *P. gingivalis* SLs delivered by OMVs have an important role in the regulation of that process. We suggest that *P. gingivalis* OMVs initiate pathogen recognition and/or inflammatory signaling in a way that can be limited by its synthesis of certain subtypes of SLs. While the molecular mechanisms utilized by *P. gingivalis* to exist in periodontal health and in disease remain incompletely defined, this duality requires further investigations to better understand how this organism utilizes SL-containing OMVs to manipulate the host-pathogen interaction.

## MATERIALS AND METHODS

### Bacterial growth and conditions

For these studies we used wild-type (WT) *P. gingivalis* strain W83, and a previously characterized isogenic mutant incapable of producing SLs due to deletion of PG1780 gene encoding the enzyme serine palmitoyl transferase (W83 ΔPG1780; SPT-) (31). Bacteria were cultivated on blood agar plates (BAPHK), and in Trypticase Soy Broth (TSBHK) at 37°C in an anaerobe chamber. Bacteria were harvested from broth culture by centrifugation, washed three-times with RPMI-1640 medium adjusted to approximately 4×10^9^ CFU/ml and were added to the upper well of the transwell cell culture system to achieve ratio of 2000 bacteria per host cell.

### Cell culture

The human monocyte cell line THP-1 (ATCC®, TIB-202, Manassas, VA) was cultured in RPMI-1640 (Corning) supplemented with 10% heat-inactivated fetal bovine serum at 37°C in a 5% CO_2_ incubator as we have done previously (17). THP-1 cells were adjusted to 5×10^5^ viable cells/ml and placed into fresh medium with 100 ng/ml phorbol 12-myristate 13-acetate (PMA; Sigma-Aldrich, St. Louis, MO) to induce differentiation into a macrophage-like state. After 48 hrs incubation, and cell washing, sterile 0.4 μm transwell inserts were placed in the wells with THP-1 cells and the inserts were filled with 125 μl of medium, *P. gingivalis* W83 parent strain or SPT-mutant as described above. After 2, 6 and 24 hrs, the cell culture supernatant fluids were collected and the levels of TNF-α, IL-1β, IL-6, IL-8, IL-10 and RANTES were determined by Milliplex Multiplex Assays (EMD, Millipore, Billerica, MA). Data was acquired on a Luminex 200® system running xPONENT® 3.1 software (Luminex, Austin, TX) and analyzed using a 5-parameter logistic spline-curve fitting method using Milliplex®Analyst V5.1 software (Vigene Tech, Carlisle, MA).

### *P. gingivalis* OMV purification and characterization

WT *P. gingivalis* W83 and the SPT-mutant were cultured in broth media as described above and OMVs were isolated and characterized as previously described (19) with modification. Bacteria were grown initially in TSBHK medium before being grown in RPMI medium under anaerobic conditions at a final volume of 500 ml. The cultures were moved to an aerobic incubator for 6 hrs prior to harvesting, to mimic conditions of the transwell experiments. Cultures were clarified by centrifugation at 7000 xg for 30 min. The supernatant was filter-sterilized (0.2 μm pore size PES membrane) then clarified supernatants were concentrated by ultrafiltration at 40°C using a Millipore stirred ultrafiltration apparatus (Millipore-Sigma; Burlington, MA). Concentrated supernatants were then ultracentrifuged at 100,000 xg for 2 hrs at 40°C to pellet crude OMVs, and the pellets were resuspended in 45% OptiPrepTM (Sigma-Aldrich) density gradient medium in HEPES buffer and overlaid with a continuous OptiPrepTM density gradient (45%-15%) and centrifuged at 100,000 xg for 16 hrs. Fraction(s) containing the OMVs were suspended in HEPES buffer then centrifuged at 100,000 xg for 2 hrs. The resultant purified OMV pellet was resuspended in a minimal volume of HEPES. Particle enumeration and size distribution were determined by NanoSight NS300 (Malvern Panalytical; Malvern, UK). The concentration of OMVs were normalized and were added directly to THP-1 cells at a ratio of 1000 particles per host cell, as previously described (51).

### Gingipain activity assays

The activity of arginine and lysine gingipains was assessed as previously described (31) (52). Cultures were inoculated from a plate into TSBHK, grown for 24 hrs then diluted into RPMI for experimental samples. Cultures were grown to exponential phase, normalized to OD_600_ = 1.0, and 1 ml of each culture was centrifuged. The supernatant fluids of these cultures were saved and assayed, and the pellets were resuspended in assay buffer (200 mM Tris, 5 mM CaCl2, 150 mM NaCl, either in the presence or absence of 10 mM L-cysteine at pH 7.6). Cultures and supernatants were diluted 1/10 in assay buffer and then serially diluted in assay buffer across a 96-well microtiter plate. The initial optical density (OD405) was recorded, and the plates were placed in a 37°C incubator for 10 min to equilibrate the temperature. N-α-benzoyl-L-arginine-p-nitroanilide (BAPNA) or N-α-acetyl-L-lysine-p-nitroanilide (ALPNA) were added to the wells at a final concentration of 1 mM, and the microtiter plates were incubated for 2 hrs at 37°C. The final optical density of the wells was recorded, and the difference between the initial and final optical density was reported.

### Quantitative real time PCR

Total RNA was extracted from cell lysates using RNAeasy kit, (Qiagen, Germantown, MD) according to the manufacturer’s protocols. The quantity of total RNA was determined by UV spectrophotometry as well as the purity, by the 260/280 nm ratio. For each sample, 500 ng of total RNA was converted into cDNA using a High Capacity cDNA Synthesis kit (Applied Biosystems, Foster City, CA). The qPCR reactions were performed in a 20 μL total volume reaction, utilizing TaqMan qPCR master mix (TaqMan Fast Advanced, ThermoFisher, Waltham, MA), cDNA template, deionized water, and human-specific pre-designed primers and probe for TLR-2, TLR4, MyD88, TICAM/TRIF, and the housekeeping gene β-actin (TaqMan gene expression assays, ThermoFisher). Cycling conditions were pre-optimized by the supplier, and 40 cycles were run on a StepOne Plus qPCR thermocycler (Applied Biosystems). Relative levels of gene expression were determined by the ΔΔCt method using the thermocycler’s software and automated detection of the cycle threshold (Ct). Expression levels of β-actin in the same samples were used to normalize results.

### Statistical analysis

Data were obtained from at least three independent experiments. Immunoassay and gene expression data were collected and both descriptive and comparative statistical analysis were performed using GraphPad 8.0 (GraphPad; San Diego, CA), non-paired t-test with Welch’s correction for unequal variances, and the significance level was set at 95% (p < 0.05) for all analyses.

## ACKNOWLEDGMENTS

We would like to acknowledge Ms. Marnelli Tan for assisting with OMV immunoassays, and the UF ICBR for use of the NanoSight. All authors acknowledge and agree to their contributions as authors on these studies and agree to its content and authorship order. These studies were supported by PHS/NIDCR grants R01DE024580 and R01DE019117 (MED), and UF Start-up (MED and FCG).

---

**Supplemental Figure 1.**
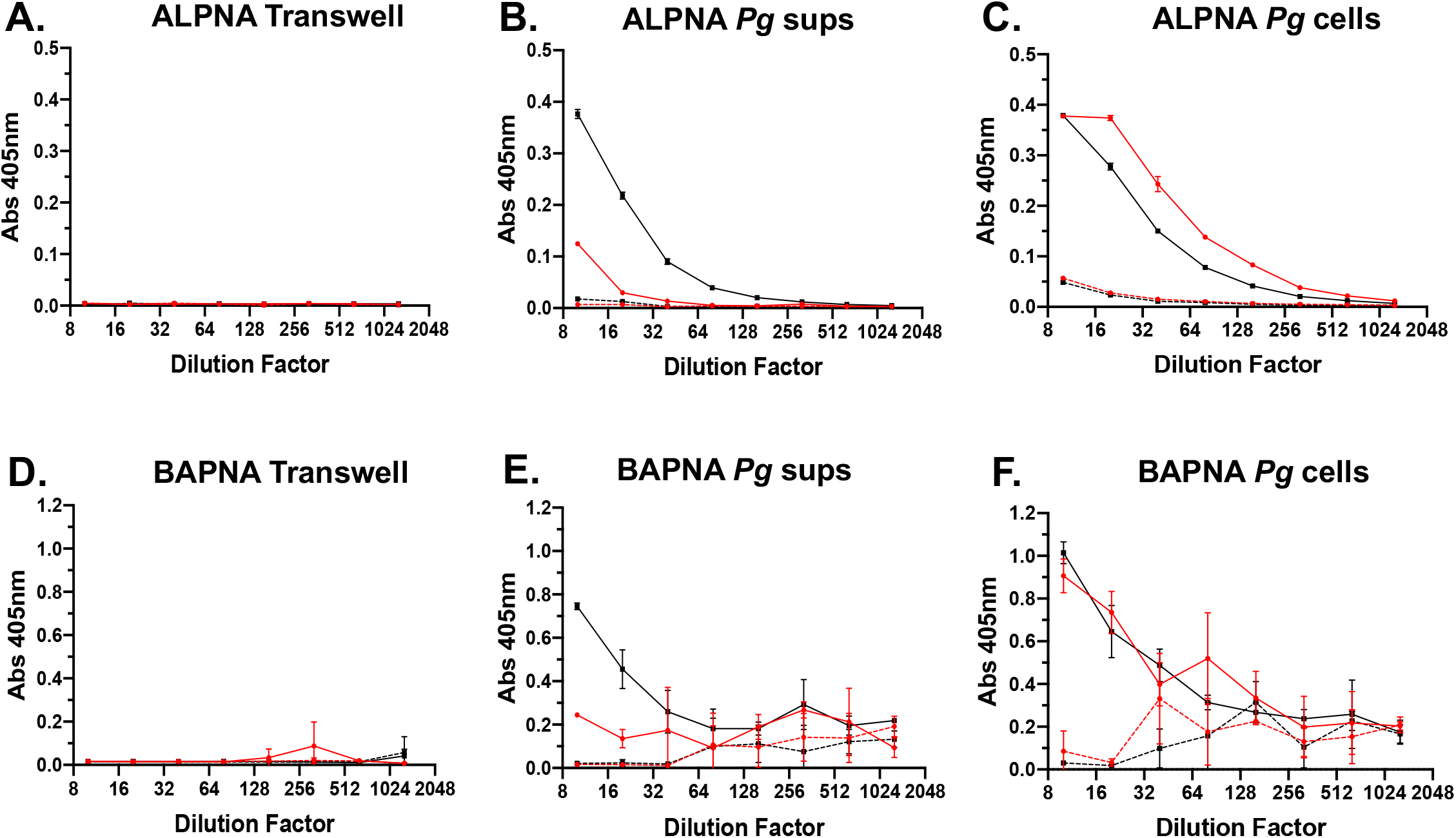
Gingipain activity present in transwell system, as we all *P. gingivalis* culture supernatant fluid and cell pellet. Culture supernatant fluids were collected from the bottom chamber of transwell experiments (**A.** and **D.**), as well as from both the bacterial cell culture supernatant fluids after separation from bacterial p>cells by centrifugation (**B.** and **E.**) and from the accompanying *P. gingivalis* cell pellets (**C.** and **F.**) these samples were assayed for lysine gingipain (ALPNA; **A.**, **B., and C.**) and arginine gingipain (BAPNA; **D., E., and F.**) activity. Transwell culture supernatant fluid levels for gingipain activity were low for W83 (red traces) and SPT-mutant (black traces) whether exogenous L-cysteine was absent (dashed lines) or present (solid lines) in the assay medium. Culture fluids collected from *P. gingivalis* following growth in TSBK, and incubation in RPMI-1640 under cell culture conditions in the absence of host cells for 6 hrs were found to have low gingipain activity in the absence of supplemental L-cysteine addition; however, it was noted in both arginine- and lysine-gingipain activity was highest in culture supernatants of SPT-mutant compared with WT *P. gingivalis.* Cell pellets of *P. gingivalis* corresponding to these bacterial supernates following 6 hrs incubation in RPMI-1640 were found to have robust arg and lys gingipain activity with W83 (red trace) possessing elevated lys-gingipain activity compared with SPT-mutant (black trace) yet possessed similar arg-gingipain activity to SPT-when assayed in supplemental L-cysteine (solid lines). Low gingipain activity was detected in the absence of supplemental L-cysteine addition (dashed lines). Data presented as mean +/− SD for n=3 separate measurements.

